# Semantic networks as a tool for analyzing conceptual organization in active teaching methodologies in Microbiology

**DOI:** 10.64898/2026.08.18.745559

**Authors:** Carolina Diorio Nastaro, Bruna Rodrigues Corrêa, Giovana Tarantini, Sandro Roberto Marana, Rita de Cássia Café Ferreira

**Affiliations:** Institute of Biomedical Sciences, University of São Paulo, São Paulo, Brazil; Institute of Chemistry, University of São Paulo, São Paulo, Brazil

## Abstract

Active teaching methodologies have been widely used to promote meaningful learning and student autonomy. In this context, quantitative approaches can help assess how students organize and integrate knowledge throughout the learning process. Among these approaches, semantic co-occurrence networks stand out, as they are capable of identifying relationships between words and revealing the conceptual structure of textual productions. The objective of this study was to investigate whether semantic network analyses can characterize differences in students’ conceptual organization in Microbiology during their participation in the active teaching methodology “Adopt a Bacterium.” To this end, a case study was conducted in the Bacteriology course at the Institute of Biomedical Sciences of the University of São Paulo, analyzing the textual productions of two groups of students in the years 2024 and 2025 during their study of the bacterial genus *Bacillus*. The texts were evaluated using semantic co-occurrence networks, taking into account metrics of structure and conceptual integration. The results showed that both groups covered the microbiological content outlined in the course, though with different thematic focuses and approaches to integrating the concepts. Although both years featured modular structures (a statistical mode of 9 subgraphs), in 2025 the network exhibited greater discursive robustness (2 to 4 times more words with high Betweenness centrality) than in 2024. It is concluded that semantic network analysis allows for the characterization of differences in conceptual organization among students using active learning methodologies, serving as a complementary tool for assessing meaningful learning in Microbiology.

## Introduction

The importance of a high quality education is widely acknowledged, hence a variety of hypotheses and methodologies have been developed to foster effective learning. To evaluate the effectiveness of these educational approaches, both qualitative and quantitative methods can be employed to assess how well they facilitate the acquisition and retention of knowledge. Quantitative research in education is of particular interest, as it allows for an objective analysis of reality and the generalization of the results obtained. Furthermore, technological resources such as software and spreadsheets support the analysis, interpretation, and presentation of data and results. [1].

Among the quantitative tools that can be used are semantic co-occurrence networks, which analyze textual data by generating graphs that show words that consistently appear close to one another, revealing the relationships between words within a message, defining the strength of these connections, and identifying themes and topics in unstructured data [2, 3]. In this way, a map of the text is generated, in which it is possible to locate clusters of words related to the theoretical topics of interest to the researcher [4], allowing for great versatility in the application of this method, which can be used in various contexts and for different subjects.

Among the types of written texts to be analyzed are those produced by undergraduate students during their coursework and while writing papers. One particular example is the Project #Adopt, an umbrella term for an initiative based in the Department of Microbiology at the Institute of Biomedical Sciences of the University of São Paulo (USP), which promotes the teaching of Microbiology to audiences of various age groups through a variety of dynamic approaches and active teaching methodologies. For this study, we highlight two of the activities developed by #Adopt: “Adopt a Bacterium” [5–7] and the production of science communication materials [8, 9].

Undergraduate students in the Biomedical Sciences program at USP take the required course Bacteriology, in which the active teaching methodology “Adopt a Bacterium” is applied. Students are divided into groups, and each group is responsible for a specific bacterium or bacterial genus. Throughout the course, students must create three posts for the social media platform Instagram®, in which they apply the theoretical knowledge covered in class to their specific bacterium, developing research skills, the ability to present collected data, and autonomy [10, 11]. Upon completing the three posts — that is, after finishing the “Adopt a Bacterium” methodology, but still within the Bacteriology course — students must submit a final project in the form of a science communication material about the bacterium or bacterial genus they worked with during the course; the format of this material (pamphlet, board game, booklet, musical parody, among others) is left to the students discretion.

In 2024 and 2025, one of the bacterial genera selected as part of the “Adopt a Bacterium” initiative was the genus *Bacillus*, a group of spore-forming bacteria of great medical importance (including the pathogenic species *B. anthracis* and *B. cereus*), as well as pharmaceutical and industrial significance (due to their widely studied applications, such as human and animal probiotics, production of enzymes and antibiotics), biotechnological (expression of recombinant proteins, use of endospores as a vaccine platforms, among others), and biomedical (*B. subtilis* strains are widely used as a model bacterium species in research), which highlights the importance of studying it by students in the Biomedical Sciences program [12, 13]. The student groups were also responsible for preparing science communication materials regarding the genus *Bacillus*.

Indeed, analyzing the content produced by these students, across both years and activities, using semantic networks can reveal how they internalized, organized, and interrelated the theory they learned in Microbiology, as well as investigate which theoretical framework each group adopted. Given the pedagogical context of active teaching methodologies, in which student autonomy is one of the key aspects [11], it is expected that different frameworks [14] and levels of interconnection will be observed as a reflection of meaningful learning.

## Materials and methods

### Participants and educational context

We analyzed the texts produced by two groups of undergraduate students who studied the bacterial genus *Bacillus* during the Bacteriology course at the Institute of Biomedical Sciences of the University of São Paulo (ICB-USP). One group was from 2024 (n = 9; women/men = 3/6) and the other from 2025 (n = 9; women/men = 5/4). All participants were undergraduate students in the Bachelor’s program in Biomedical Sciences in their second semester of the program. In both years, the Bacteriology course, offered by the Department of Microbiology, was taught by the same instructors, using the same teaching materials, teaching methodology, and theoretical content.

### Instructional design

The Bacteriology course uses the active teaching methodology “Adopt a Bacterium” [11, 5–7] as a pedagogical tool. In this approach, Undergraduate students are organized into groups, each assigned a bacterium or bacterial genus to study (“adopt”) throughout the semester. After attending in-person lectures, students apply the knowledge they have acquired to their assigned microorganism, taking responsibility for researching the information and presenting it in the form of posts (n = 3) on the social media platform Instagram®.

The content covered in class, which was required to be addressed in the posts, was the same in both years: morphology, diversity and ecology, nutrition, growth and metabolism, biochemical tests, genetics, antimicrobials, microbiota, growth control (chemical and physical agents), and pathogenicity. Students were free to include additional information, provided that the required content was included. The presentation of this information, that is, the way they choose to convey their research (using illustrations, comic strips, personification, storytelling, among others), is left at the group discretion.

At the end of the course, both groups submit a project in the form of a science communication material based on all the research conducted during “Adopt a Bacterium.” The type of science communication material, such as pamphlets, games, musical parodies, magazines, and others, is also at the group discretion.

### Texts written by the students

In 2024 (S1 Fig), the group chose to use a storytelling approach for their social media posts, featuring the characters “Wellington the Mouse,” his daughter “Wanda the Mouse,” and the doctor “Wilson the Mouse” as a narrative thread to present their research. As for the science communication material, the 2024 group created a magazine that combined theoretical content with variety puzzles, such as word searches and crosswords. The group stated that the target audience for the material was high school students.

In 2025 (S2 Fig), the group also used storytelling to organize its social media posts, which were presented in a blog format. In addition, they used the personification of bacterial species (*Bacillus subtilis* as the influencer “Bah,” author of the blog; *B. cereus* as “Cousin Cici,” among others) as a strategy for presenting the content. As for the science communication material, the group developed a card game in which participants must solve mysteries based on their knowledge of *Bacillus* characteristics. The group stated that the target audience for the material was high school students or those in their early years of college, though not necessarily in a health-related field.

The texts of the posts and science communication materials were analyzed based on the number of sentences, the number of paragraphs, the average number of words per sentence, and the TTR (Type-Token Ratio) parameter, a measure that indicates the lexical variety of the text by calculating the ratio of the number of distinct words (Types) to the total number of words (Tokens) [15].

### Analysis software

Textual analysis, including text processing and graph construction, was performed using the KH Coder application (https://khcoder.net/en/), an “open-source software program, distributed free of charge, released in 2006 by Koichi Higuchi for computational linguistics or quantitative content analysis” [16]. KH Coder also offers a wide range of analytical tools and support for Portuguese-language texts. For this study, the free version of the software was used.

### Text preprocessing

For the analyses, only texts containing full-length written content were considered (social media captions and text contained in images were disregarded). In the case of science communication material, only the text was considered, disregarding the words contained in the instruction manual (for the 2025 group) and the words in the variety puzzles (for the 2024 group).

The texts were tabulated in Microsoft Excel®, and the content was spell-checked. Abbreviated words were rewritten in their full form (for example, replacing the abbreviation “*B.*” with its full form “*Bacillus*”). Data normalization was performed directly and automatically by the semantic analysis software KH Coder, which involved the removal of stop words (words such as articles, pronouns, and prepositions that occur frequently but lack relevant semantic information) and lemmatization of the words (reducing a word to its root form by removing number and conjugation inflections).

### Construction of semantic networks

The co-occurrence semantic networks [2] were constructed directly in the free version of the KH Coder application. The networks, also called graphs, show correlated words (nodes) connected by an edge, thus representing the fact that they have similar patterns of occurrence [3]. The size of the node is proportional to the frequency with which the word appears in the text. In KH Coder, words that were correlated but whose co-occurrence was not strong enough to be plotted on the graph are also displayed. In this case, these words are shown as white nodes, and the weak correlation is indicated by dashed edges.

The correlation between words is calculated using the Jaccard coefficient [17], which assesses the number of times any two words appear close to each other (words are considered “close” if they appear within 10 words of each other) relative to the total number of times they appear in the text. The graph is plotted in space using the Fruchterman-Reingold algorithm [18]. Consequently, it should be noted that in the graph, only words or terms connected by an edge are correlated, whereas their spatial proximity is not relevant to the analysis.

### Network analysis metrics

The metrics used to interpret the data were the number of nodes (representing lexical diversity), the number of edges (representing the number of conceptual relationships), and network density (the ratio of the number of edges to the total possible number of edges, that is, conceptual relationships that occurred relative to all possible relationships). In this sense, higher density is associated with greater conceptual interconnectedness in the group texts.

Graph modularity was also used as a metric to indicate conceptual segmentation or integration. It was calculated using the modularity maximization algorithm developed by Clauset *et al*. [19], implemented by the KH Coder software.

Finally, Betweenness centrality was used to identify terms that act as bridges in the word co-occurrence network. According to the formulation by Newman & Girvan [20], nodes with high Betweenness centrality tend to play a role in the structural organization of the text by connecting distinct modules. Thus, words with high Betweenness values were interpreted as elements responsible for linking different thematic topics.

### Definition of cut-off values

The cut-off values were selected based on the work of Sarachuk [16], who uses the Term Frequency (TF) distribution as an indicator of the cut-off line for minimum TF. The cut-off value was adjusted across the datasets to maintain comparability of the lexical coverage of the analysis among the materials, covering 8.74% to 9.73% of the total vocabulary in all four text corpora analyzed. The cut-off for the Jaccard coefficient was 0.3 for the posts and 0.5 for the science communication materials (Table 1).

**Table 1.** Minimum TF cut-off scores and Jaccard coefficients used for the 2024 and 2025 data.

|  | 2024 | 2025 |
| --- | --- | --- |
| **Instagram® posts** |  |  |
| Minimum Term Frequency (TF) | 3 | 4 |
| Percentage of words included | 9,55% | 9,09% |
| Number of words included | 71 | 110 |
| Jaccard coefficient | 0.3 | 0.3 |
| Density of the resulting co-occurrence graph | 0.09 | 0.039 |
| **Science communication materials** |  |  |
| Minimum term frequency (TF) | 2 | 3 |
| Percentage of words included | 9,73% | 8,74% |
| Number of words included | 51 | 105 |
| Jaccard coefficient | 0.5 | 0.5 |
| Density of the resulting co-occurrence graph | 0.129 | 0.029 |

Minimum Term Frequency (TF): the minimum number of times a word must be present in the text in order to be included in the analysis, adjusted so the percentage of words included in each case was similar. Jaccard coefficient: the minimum correlation strength between two words in order for them to be included on the analysis.

### Ethics

This project (CEP ICB-USP Protocol #990/2018) was evaluated by the Research Ethics Committee of Human Beings (CEPSH ICB-USP) and was considered exempt from the need for consent form (report #1247 issued on 11/26/2018). This exemption was because the present study did not carry out any procedures regulated by CONEP resolution #466/2012.

## Results

### Structural linguistic analysis of social media posts and science communication materials from the “Adopt a Bacterium” activity in 2024 and 2025

In 2024, the group of undergraduate students produced social media posts about *Bacillus* using the storytelling method. Those posts had a TTR (Type-Token Ratio; an indicator of lexical variety) of 0.283 and an average of 21.3 words per sentence. As for the science communication material, this group produced a variety word puzzle magazine, which had a TTR of 0.318 and an average of 17.1 words per sentence (Table 2). Similarly, in 2025, the group of students also used storytelling to organize their social media posts, which were presented in blog format. These posts had a TTR of 0.220 and an average of 23.4 words per sentence. As for the science communication material, the 2025 group produced a card game, with a TTR of 0.273 and an average of 18.2 words per sentence (Table 2).

**Table 2.** Structural linguistic analysis of the texts produced in the “Adopt a Bacterium” learning activity.

|  | 2024 |  | 2025 |  |
| --- | --- | --- | --- | --- |
|  | Posts | Science communication material | Posts | Science communication material |
| **Tokens** | 1.879 | 1.029 | 4.793 | 2.647 |
| **Types** | 531 | 328 | 1.056 | 724 |
| **TTR** | 0,283 | 0,318 | 0,220 | 0,273 |
| **Number of sentences** | 88 | 60 | 205 | 145 |
| **Number of paragraphs** | 35 | 29 | 91 | 47 |
| **Average number of words/sentence** | 21,3 | 17,1 | 23,4 | 18,2 |

TTR (Type-Token Ratio) was used to indicate the lexical variety of each text, verifying their comparability. Types: number of distinct words in the text. Tokens: total number of words.

Although the texts produced by the 2025 group were longer than those produced by the 2024 group, as observed by the number of sentences and paragraphs (Table 2), the TTR ratio, as well as the average sentence length, remained similar across all four materials analyzed, indicating that they can be compared using semantic networks.

### Application of semantic co-occurrence networks in the analysis of the social media posts about *Bacillus* in 2024 and 2025

The co-occurrence semantic network of social media posts about the genus *Bacillus* produced during participation in the “Adopt a Bacterium” learning activity in 2024 displayed 9 subgraphs (word clusters) (Fig 1; marked in different colors). Out of the 71 words included in the analysis based on the TF cut-off score (Table 1), 68 met the minimum correlation strength criterion (Jaccard coefficient) to be plotted on the graph. A total of 206 correlations passed the cut-off established by the Jaccard coefficient. The graph density was 0.09, indicating that of the total possible correlations among the words in the text, 9% are represented in the graph.

**Fig 1.**
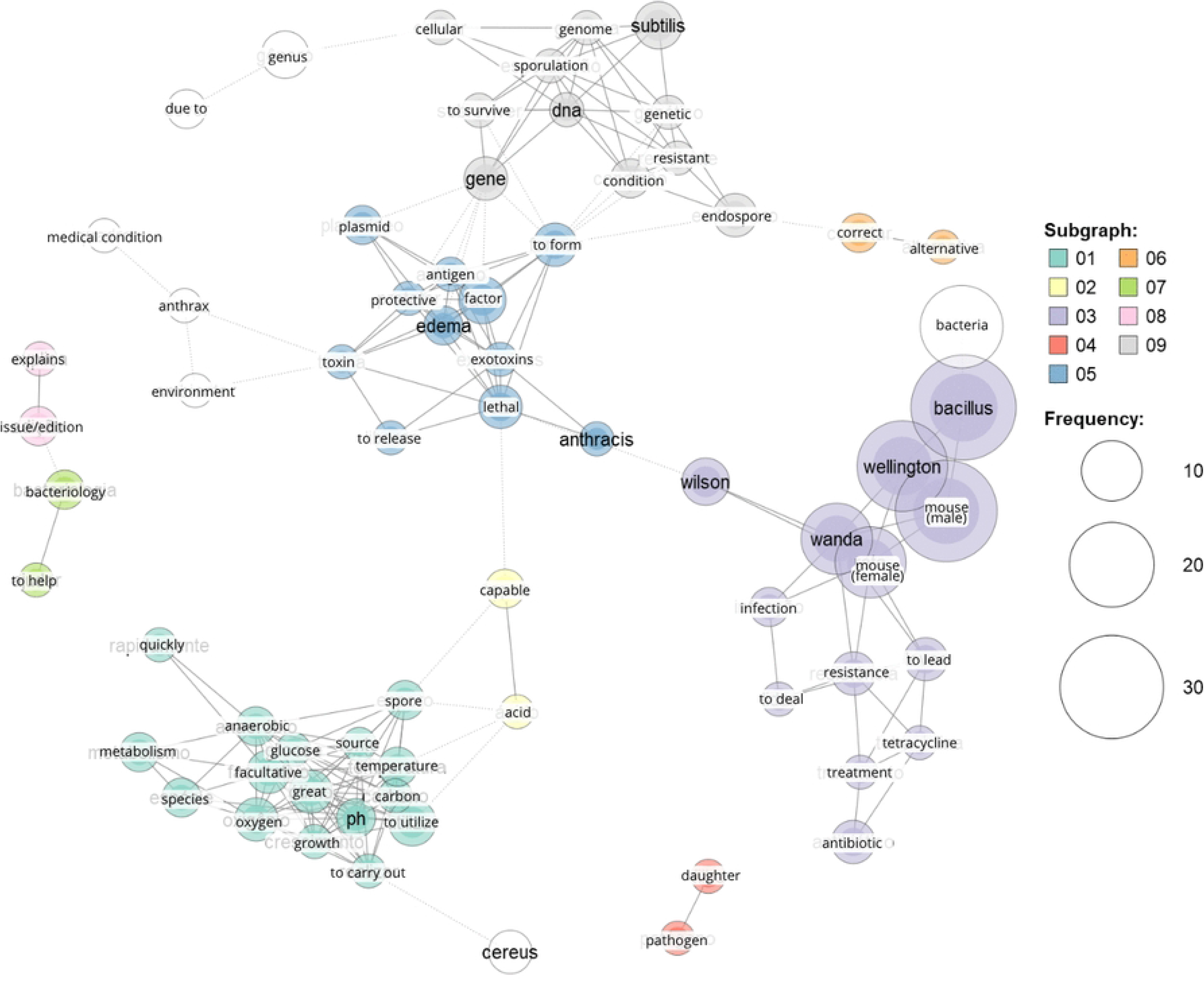
Semantic co-occurrence network for “Adopt a Bacterium,” 2024 group. Graph modularity metric indicates conceptual segmentation in 9 clusters (subgraphs), indicating low conceptual integration. Plotted nodes: 68, edges: 206, density: 0.09.

It is noticeable that the word clusters followed the Microbiology topics the groups were supposed to address in their posts, as in the subgraphs containing vocabulary related to metabolism and physiology (subgraph 1), genetics (subgraph 9), pathogenicity (subgraph 5), and antimicrobials (subgraph 3). It is also possible to identify words resulting from the content presentation strategy adopted by the group, as the character names in subgraph 3 (“Wellington the Mouse,” “Wanda the Mouse,” “Wilson the Mouse”), and the fictional scenario in which they were involved in subgraph 4 (Wanda the Mouse is Wellington the Mouse’s “daughter” and was infected by a “pathogen”). Also, words related to the context and purpose of the posts are present in subgraphs 7 and 8 (what would be “explained” in each “issue” posted on Instagram® and how to “help” the mice learn “Bacteriology”).

It is also noted that the character names appear with high frequency in the posts, on par with the word “*Bacillus*” itself (subgraph 3).

The Betweenness centrality in this network (Fig 2) revealed that words such as “lethal” (referring to the “lethal” factor and toxin produced by *Bacillus*), “capable,” “to form,” “spore,” and “Wilson” stand out. Given the definition of Betweenness centrality, we assume that these words act as “bridges”, frequently connecting different concepts within the network. Although these central words do not necessarily show strong correlations with one another (Fig 1), they were in fact used to link different conceptual subgraphs (word clusters).

**Fig 2.**
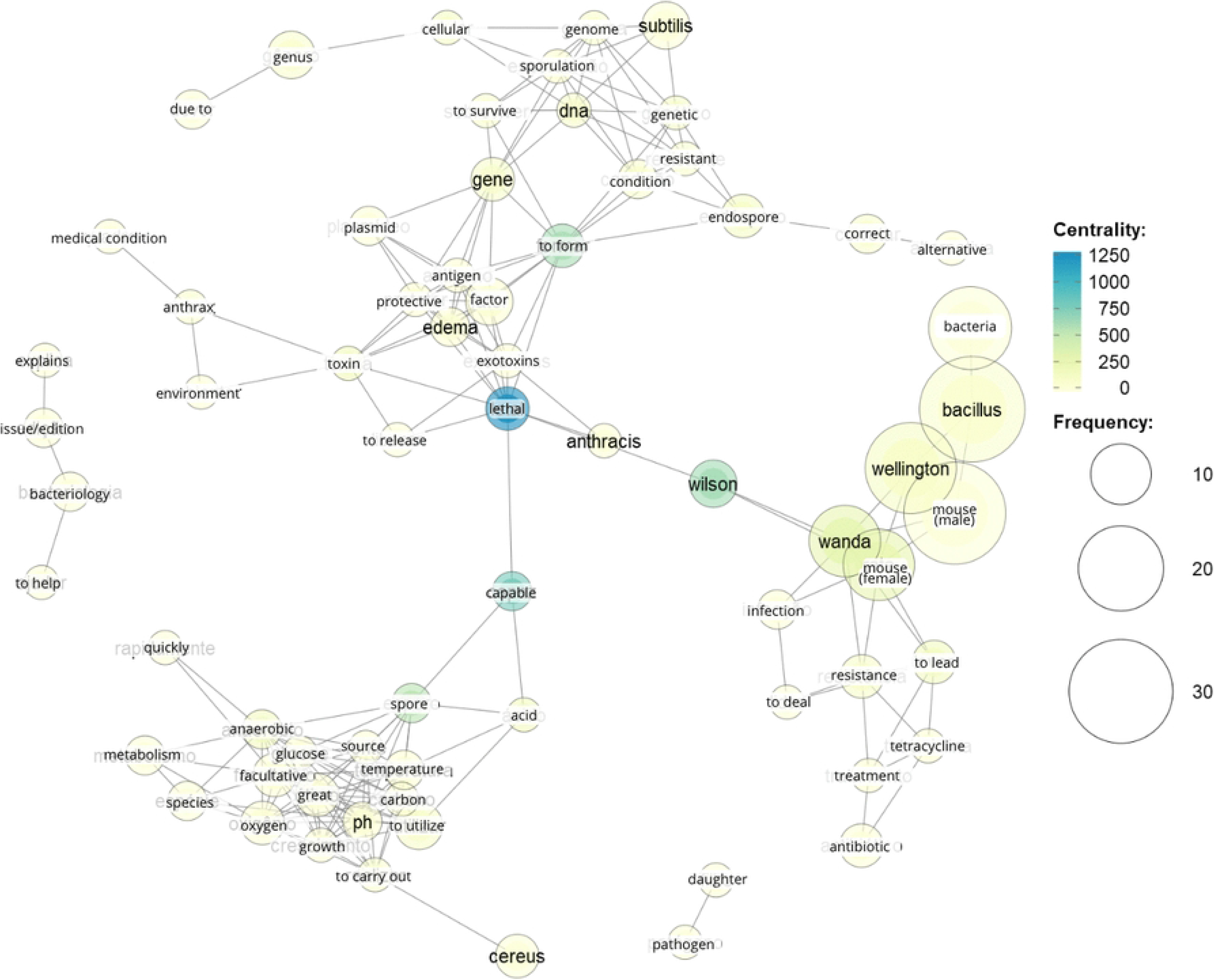
Semantic co-occurrence network for “Adopt a Bacterium,” 2024 group, showing the centrality of the words. Betweenness centrality was used to identify terms that act as bridges in the word co-occurrence network. Thus, words with high Betweenness values are responsible for linking different thematic topics. Only “lethal” exhibits high centrality (> 1000), illustrating the fragility of this thematic interconnection.

The graph resulting from the social media post in 2025 (Fig 3) showed 102 nodes and 203 edges, with a density of 0.039. Thus, in this year, the number of correlations identified (3.9%) in the social media posts was lower compared to 2024 (9%). On the other hand, as in the previous case (Fig. 1), 9 subgraphs were also detected in the network from 2025, and the words clustered within also align with the topics that students were required to cover. However, words in the 2025 network are not as clearly segregated as in the 2024 graph (Fig 1), showing terms related to toxin-mediated pathogenicity observed in both subgraph 6 (“diarrhea,” “formation” of “pores” in “cells,” non-hemolytic enterotoxin “NHE,” and hemolysin BL “HBL”) and subgraph 4 (“nausea,” “vomit,” “toxin”). Subgraph 1, on the other hand, contains information on both metabolism and physiology (“metabolism,” “pH,” “temperature,” “nutrient”) and genetics (“gene” and “DNA”).

**Fig 3.**
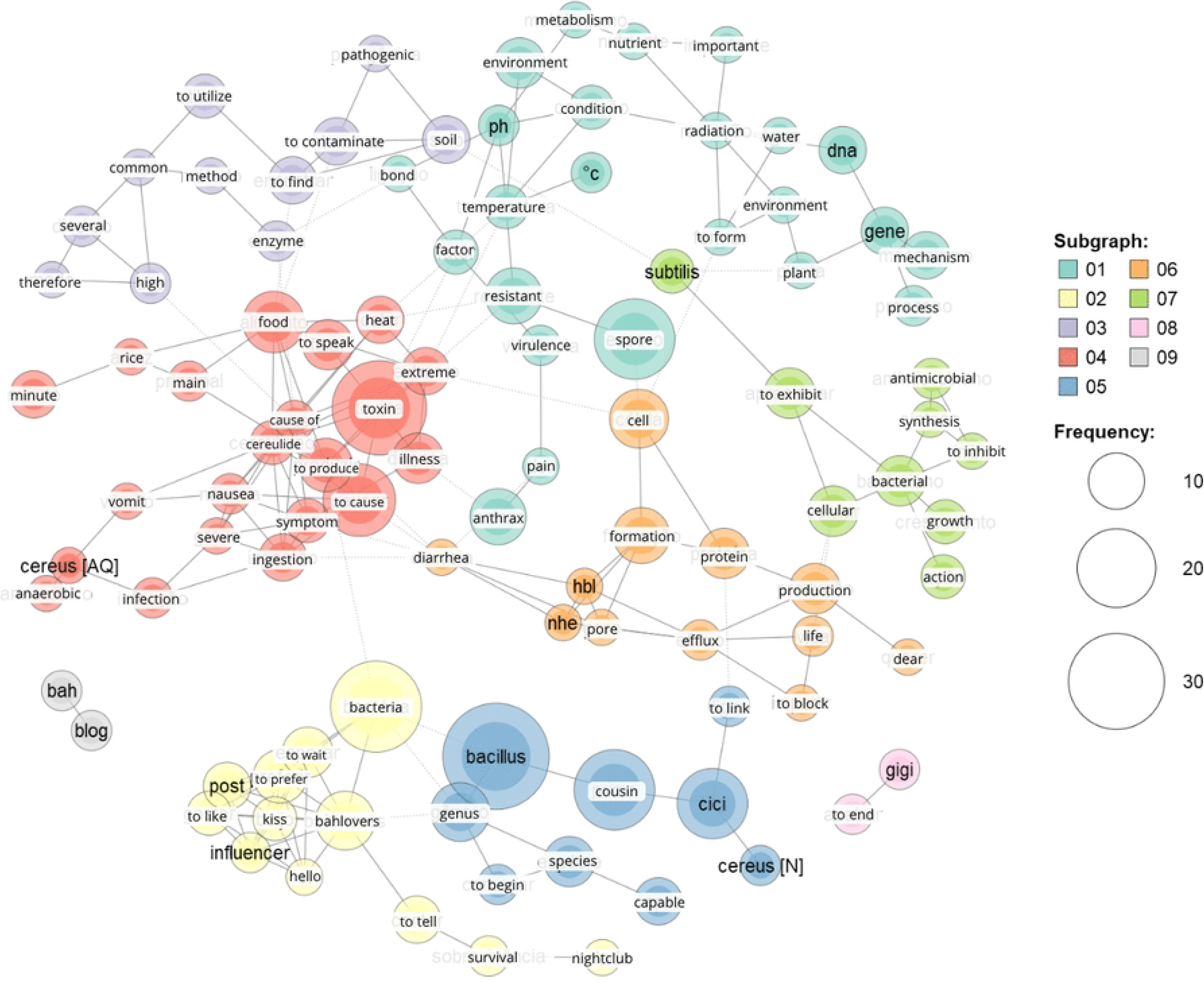
Semantic co-occurrence network for “Adopt a Bacterium,” 2025 group. Graph modularity metric indicates conceptual segmentation in 9 clusters (subgraphs), indicating low conceptual integration. Nodes plotted: 102, edges: 203, density: 0.039.

Once again, we detected subgraphs with terms that refer to the fictional setting of the characters created by the group, as observed in subgraphs 9 (“blog” of “Bah,” the name of the character *Bacillus subtilis*) and 2 (“influencer,” “post,” “kiss” and “bahlovers,” the name by which the character “Bah” refers to her followers).

The most central words according to the Betweenness centrality criterion were “bacteria,” “food,” “to produce,” and “extreme” (Fig 4). Therefore, they do not coincide with the words classified in this same category in the 2024 graph.

**Fig 4.**
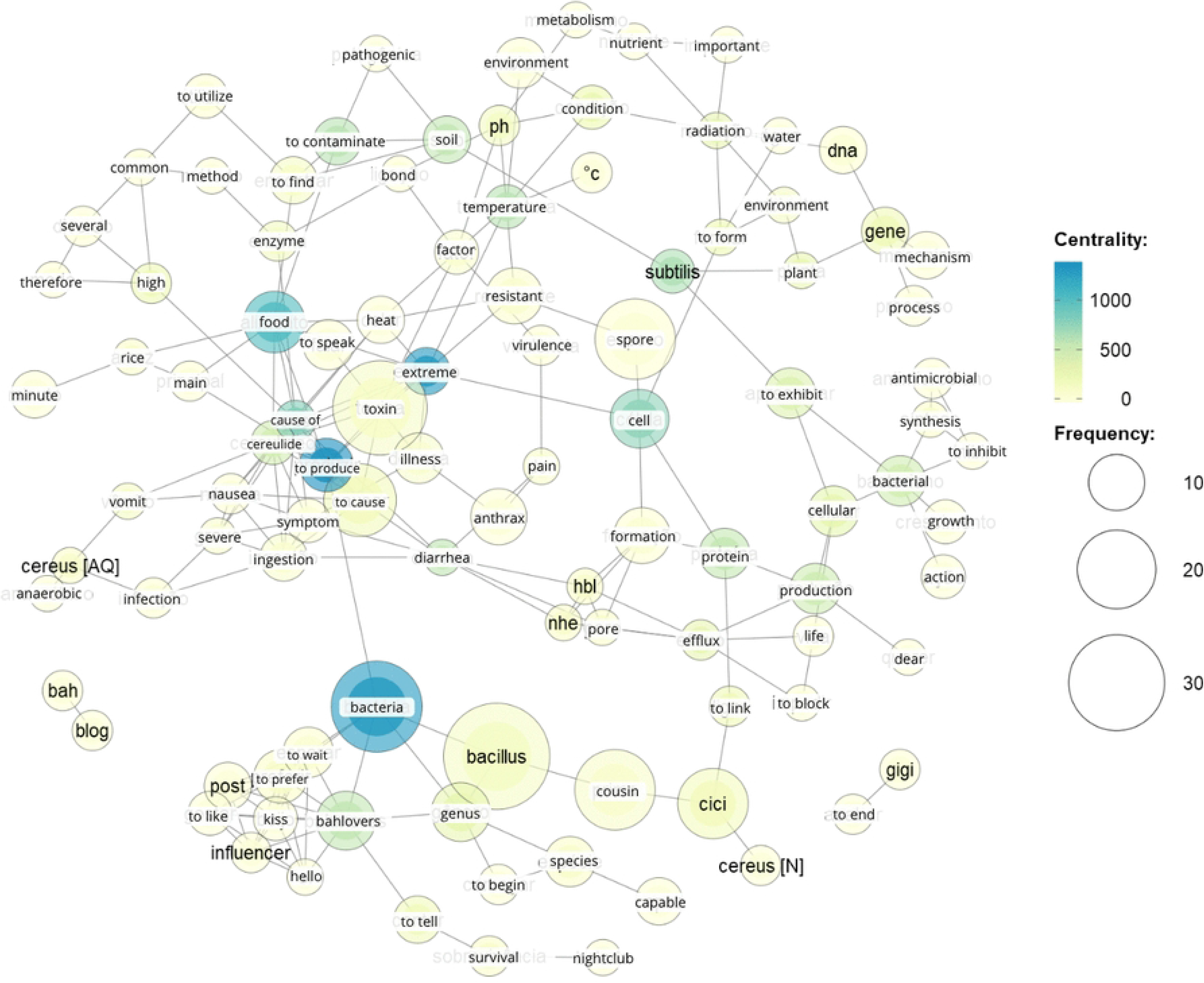
Semantic co-occurrence network for “Adopt a Bacterium,” 2025 group, showing the centrality of the words. Betweenness centrality was used to identify terms that act as bridges in the word co-occurrence network. Thus, words with high Betweenness values (“bacteria”, “food”, “to produce” and “extreme”) were interpreted as elements responsible for linking different thematic topics.

### Semantic co-occurrence network analysis of the science communication material about *Bacillus* produced in 2024 and 2025

The graph based on the science communication material produced by the 2024 group (Fig 5) consisted of 51 words and 165 correlations. It is worth noting the presence of only 5 subgraphs, whereas 9 were observed in the graph related to the posts (Fig 3), suggesting that the students chose to address a smaller number of topics in the outreach material.

**Fig 5.**
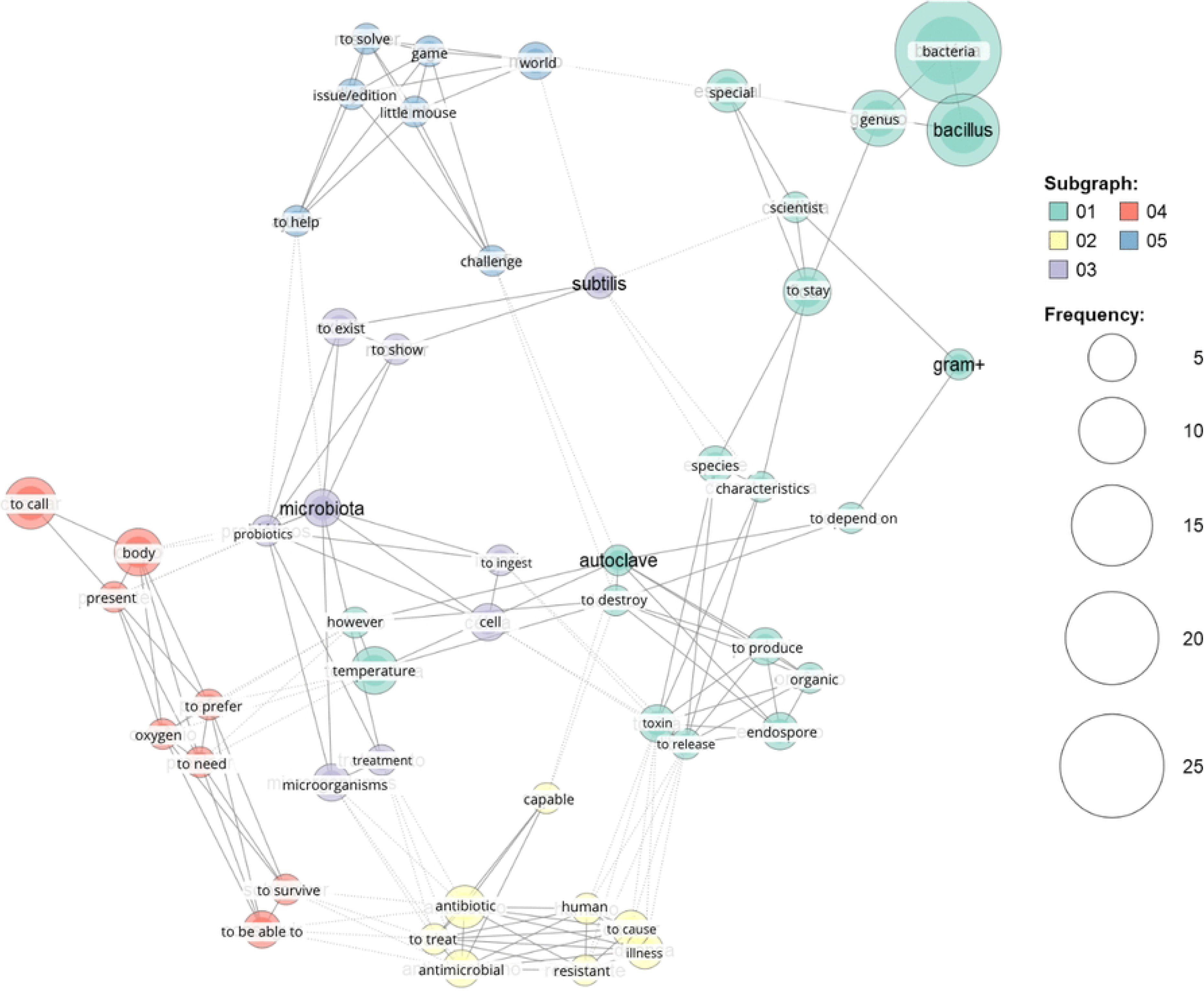
Semantic co-occurrence network of the science communication material, 2024 group. Graph modularity metric indicates conceptual segmentation in 5 clusters (subgraphs), indicating a higher conceptual integration. Nodes plotted: 51, edges: 165, density: 0.129.

**Fig 6.**
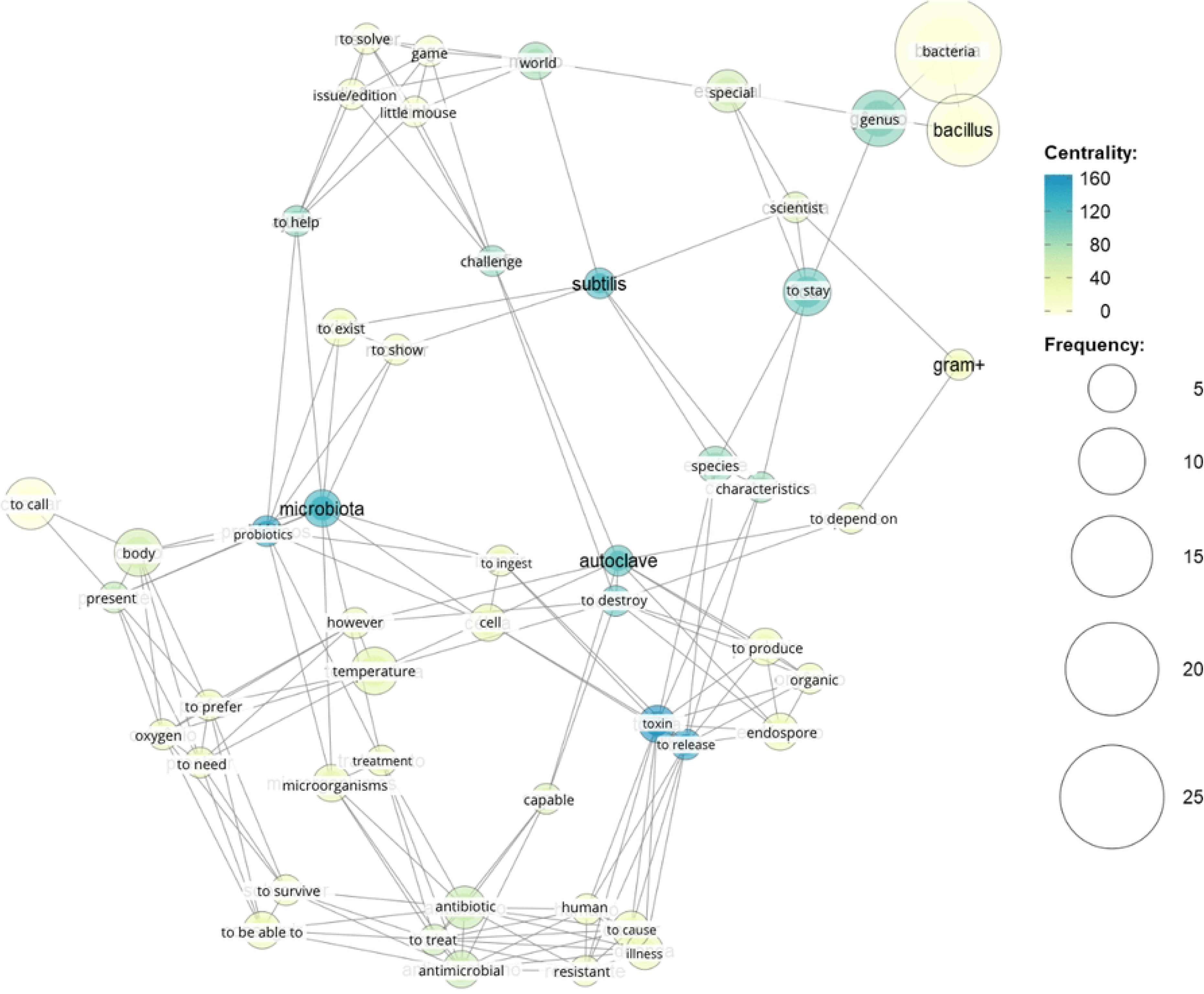
Semantic co-occurrence network of the science communication material, 2024 group, showing the centrality of the words. Betweenness centrality was used to identify terms that act as bridges in the word co-occurrence network. Thus, the low centrality seen on this network indicates relative thematic isolation.

**Fig 7.**
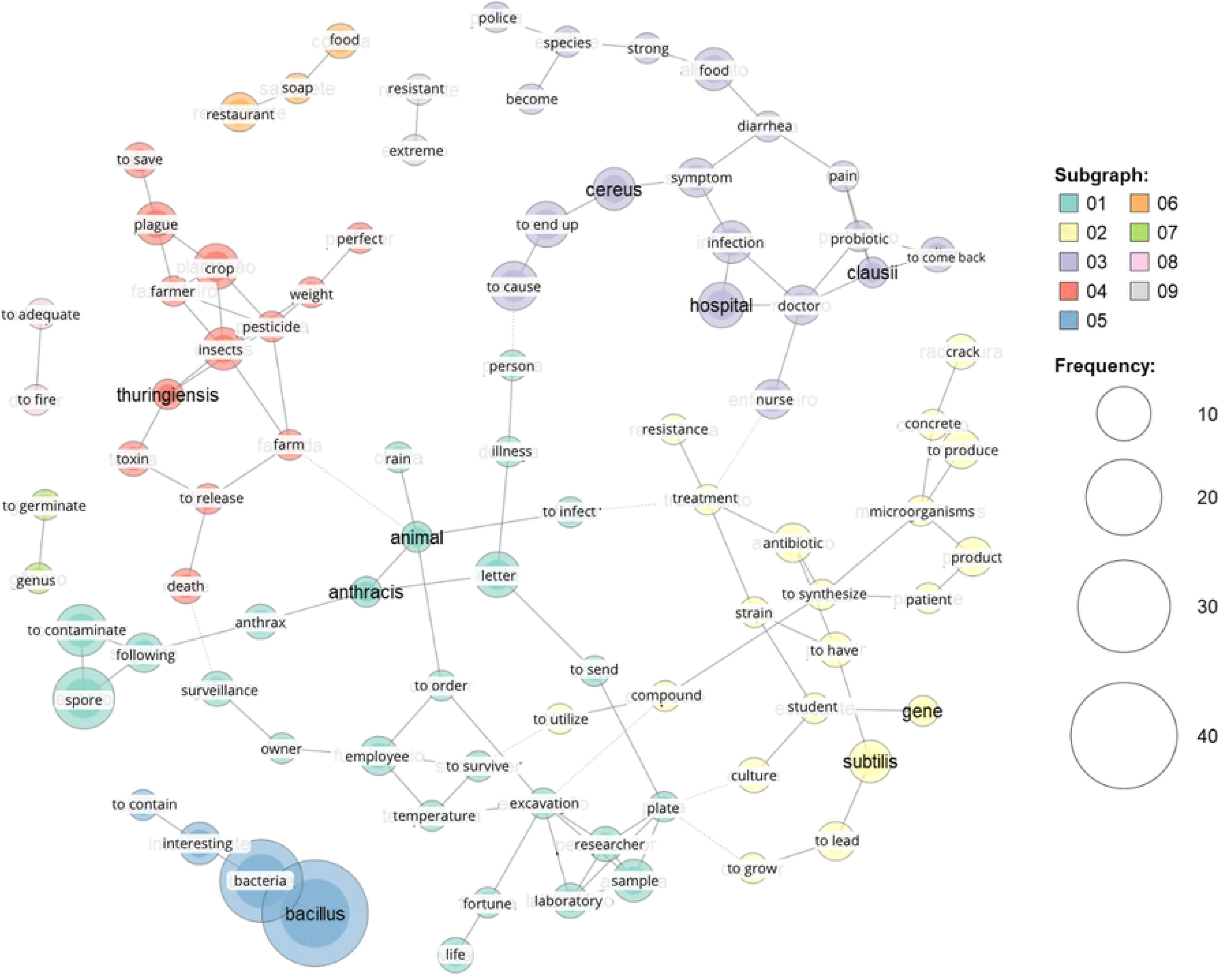
Semantic co-occurrence network of the science communication material, 2025 group. Graph modularity metric indicates conceptual segmentation in 9 clusters (subgraphs), indicating low conceptual integration. Plotted nodes: 89, edges: 113, density: 0.029.

There is a single subgraph containing words related to the context of the communication material (subgraph 5: “help” the “little mouse,” “solve” the “games” and “challenges” in this “issue” of the variety word puzzle magazine). It is observed that contextual and character-related words do not appear very frequently. In addition, the most frequently used word was “bacteria.”

The graph of this science communication material exhibits higher density compared to the social media posts graph (0.129 *versus* 0.09). However, this higher percentage of correlations did not translate into greater integration among the themes of the subgraphs, since all words exhibit low Betweenness centrality (< 250) (Fig 8). This holds true even for the central words in this network, such as “*subtilis*,” “probiotic,” “microbiota,” “toxin,” and “autoclave.”

**Fig 8.**
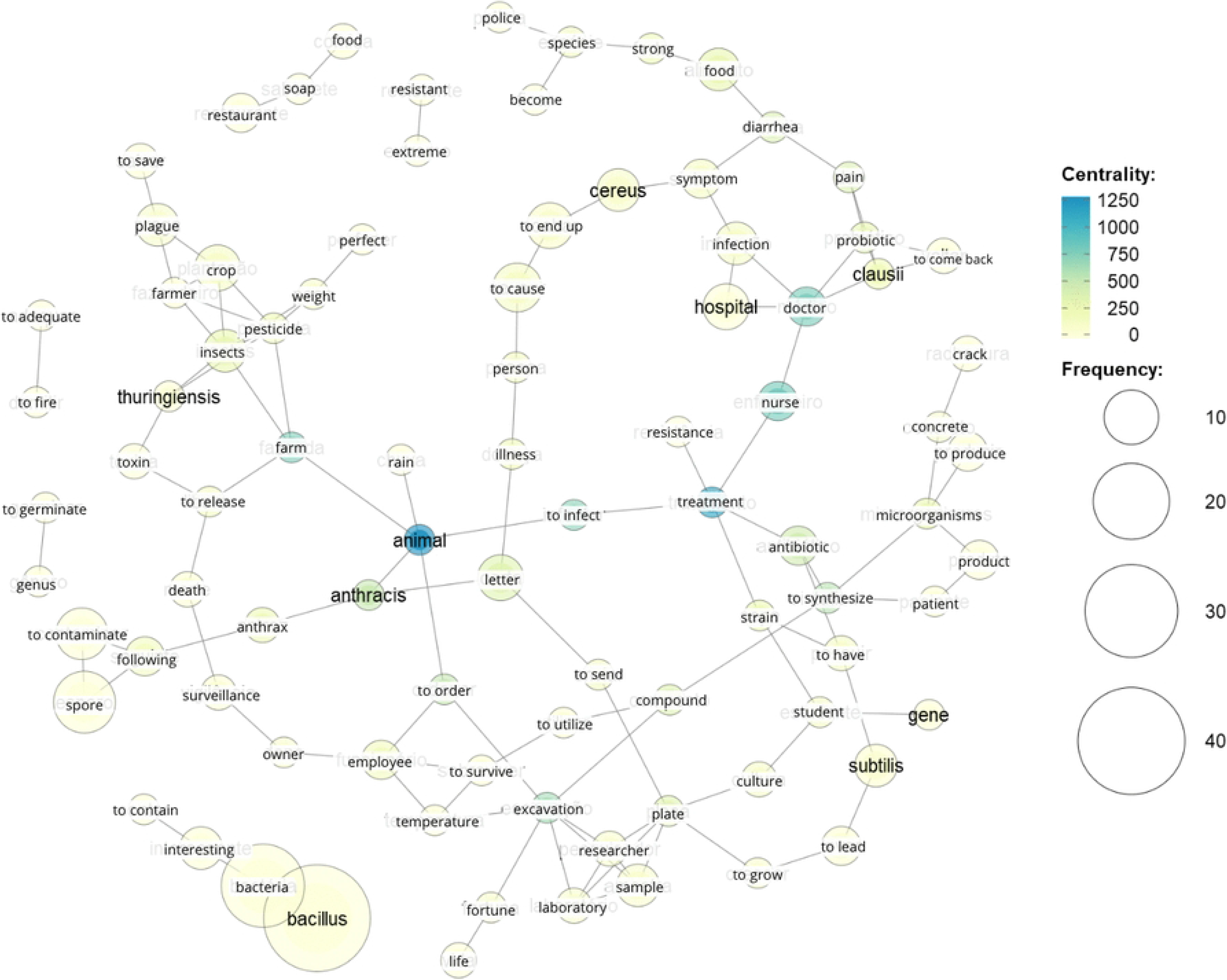
Semantic co-occurrence network of the science communication material, 2025 group, showing the centrality of the words. Betweenness centrality was used to identify terms that act as bridges in the word co-occurrence network.

**Fig 9.**
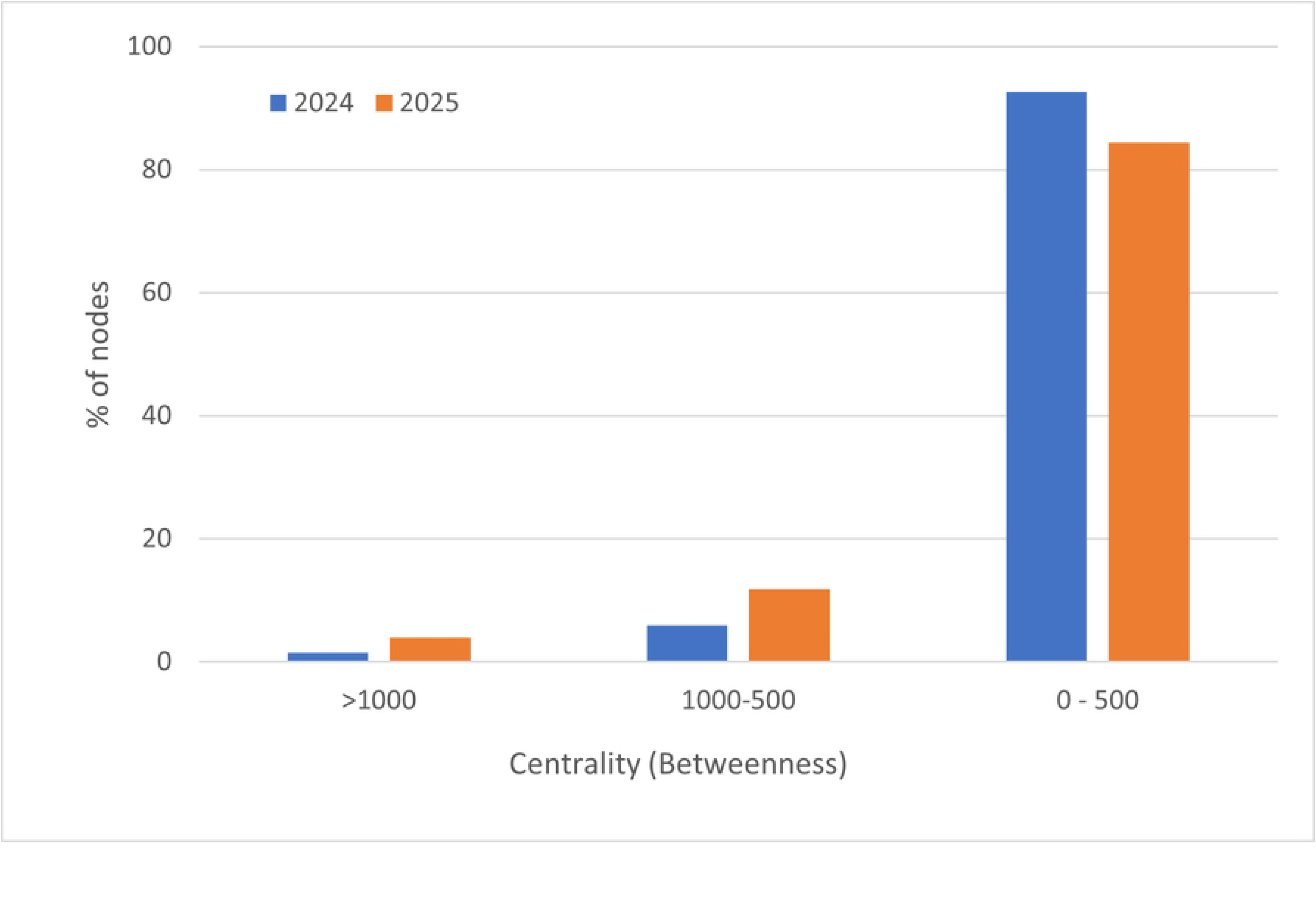
Distribution of Betweenness centrality in the semantic networks of posts produced in 2024 and 2025 during participation in the “Adopt a Bacterium” initiative. Thus, although the 2025 group also exhibited modular discourse in their social media posts, as revealed by the thematic subgraphs, the high centrality of some nodes in this network suggests a certain discursive robustness, with a greater number of words serving as bridges between all the subgraphs.

The semantic network based on the communication material produced in 2025 (Fig 7) consisted of 89 nodes (words) and 113 connections grouped into 9 subgraphs, as previously observed in the network of the social media posts (Fig 3), suggesting that all the topics studied during the course reemerged in these two contexts. It is evident that the students opted for a more biotechnological and biomedical approach in the game, as seen in subgraphs 2 (“product,” “synthesize,” “compound,” “culture,” “strain”), 3 (“probiotic,” “hospital,” “infection,” “symptom”), 4 (“pesticide,” “pest,” “crop,” “farmer”), and 1 (“sample,” “laboratory,” public health “surveillance”).

The percentage of correlations between words in the networks of social media posts (Fig 3) and science communication material (Fig 7) are similar (0.039 *versus* 0.029). Furthermore, both networks contain words with high Betweenness centrality (>1000), suggesting a similar degree of integration among the themes. The words with the highest centrality in the network of the science communication material produced in 2025 (Fig 8) were “animal,” “farm,” “treatment,” “nurse,” “doctor,” and “to infect,” themes that frequently provided context for the mysteries described in the game cards.

Regarding the Betweenness centrality of the semantic networks of the communication material, the values reached in the 2024 network fall entirely within the low centrality class, whereas 9% of the words exceed this threshold in 2025.

## Discussion

### Semantic networks and social media posts about *Bacillus* in 2024 and 2025

Semantic co-occurrence networks of words provide an overview of the topics present in text excerpts. Indeed, in the networks built based on the social media posts made by students, both in 2024 and 2025, it is clearly possible to identify subgraphs grouping terms related to the concepts that students were required to address in their research. At the same time, this tool also highlights that different groups of students have different interests, and that this is reflected in the theoretical framework they adopt and the way they present their work.

The semantic networks suggest that in 2024 the social media posts addressed the studied topics, but in a modular manner, with less integration between subjects when compared to the network from 2025. In fact, only one word (“lethal”) exhibited high centrality (> 1000) in the network from 2024, illustrating the fragility of this thematic interconnection. Further evidence of this reduced integration is that the 2024 semantic network is richer (93%) in words with low Betweenness centrality (< 500), whereas in 2025 this class of words is less abundant (84%). Consistently, in the 2025 semantic network, words with medium (500–1,000) and high (>1,000) Betweenness centrality are, respectively, 2 to 4 times more frequent than in 2024 (Fig 9).

### Semantic networks and the science communication material on *Bacillus* in 2024 and 2025

The science communication magazine prepared by the 2024 group generated a semantic network with fewer subgraphs than the number of topics studied during the “Adopt a Bacterium” program, yet with the highest observed density of intercorrelations. However, the nodes in this semantic network have low centrality. This combination suggests that the groups may have chosen specific themes from the entire sample and kept them relatively isolated, with only many internal correlations.

Meanwhile, an analysis of the 2025 communication material, a card game, revealed the same number of subgraphs and topics studied, as well as a low correlation density among words. However, this semantic network contains words with high centrality. These characteristics may stem from the way students chose to design the science communication material. Since they broke the content down into mysteries to be solved, it is understandable that they opted to reduce the overlap between the cards, while keeping only few conceptual words to mark the trail among them, in order to make the game more engaging and challenging. Thus, a low correlation density combined with high centrality words.

In this case, it is also interesting to note how the group preference for topics related to biotechnology and the practical application of *Bacillus* bacteria became evident, given that the choice of how to approach the content originated from the group itself.

This difference in the approaches and directions chosen by the 2024 and 2025 groups highlights a striking feature of active teaching methodologies: students have autonomy to develop projects on their own, since they sought out the requested theoretical content and chose for themselves how to present that content. Furthermore, the different focuses adopted by each group demonstrate Rogers educational theory, which holds that meaningful learning occurs when the content resonates with personal interests, that is, when the motivation behind that learning comes from within the student themselves.

Finally, this study highlights the role of quantitative analysis in educational research. In particular, semantic networks are capable of identifying textual topics even in non-traditionally structured datasets, as demonstrated by their ability to extract data even from card games and social media posts. It should be noted, of course, that in such cases the analysis of the results must be conducted more carefully, since it directly impacts the final network produced, as evidenced by the fact that the network derived from the 2025 science communication material exhibited low interconnection density due to the intrinsic characteristics of the proposed material style. Despite this, even in these cases, the analysis is still capable of revealing important information, such as the theoretical framework, the degree of interconnection among conceptual topics, and interests that motivate that group of students, which provides guidance for teachers seeking ways to increase student engagement in the classroom.

## Conclusions

The “Adopt a Bacterium” active teaching methodology was previously designed to combine freedom of expression and creativity with mandatory key educational components, allowing the students to learn Bacteriology while also exploring topics that interest and motivate them, a feature that enhances the chance to promote meaningful learning.

The application of the semantic networks to analyze the materials produced along the “Adopt a Bacterium” learning activity (social media posts, a magazine and a card game) confirmed that even in this informal setting the students recalled the conceptual topics as shown by the number of subgraphs within each network. In addition to that, the network analysis pointed to a different connectivity among topics depending on the year and strategy of the science divulgation material. Finally, the analysis of the subgraphs within the network revealed conceptual topics favored by students in different years, confirming their freedom to explore topics that motivate them.

The detection of subgraphs evoking Microbiology concepts also shows that the genus *Bacillus* is an excellent model for Microbiology education. Its widespread occurrence in nature, ability to form endospores, application in industry, role as a model organism in research, pathogenic species, as well as its agricultural, pharmaceutical, and biotechnological applications, open up a wide range of options for students to direct their research according to their individual interests, ensuring their motivation throughout the learning process.

In short, semantic co-occurrence networks are an important tool for quantitative analysis in education, as they can highlight the main topics addressed by students, the degree of interconnection among topics, and the interests that motivate the students, allowing teachers to tailor their lessons to generate the greatest possible student engagement and motivation.

## Acknowledgments

We thank all the undergraduate students of the Institute of Biomedical Sciences at the University of São Paulo for participating in the project.

## Supporting information

**S1 Fig. Content produced by the students in 2024. (A)** Cover images of Instagram® posts made during the “Adopt a Bacteria” project. **(B)** Cover of the magazine produced by the students as science communication material.

**S2 Fig. Content created by the students in the class of 2025. (A)** Cover images of Instagram® posts made during the “Adopt a Bacterium” project. **(B)** Front and back of two cards from the game designed by the students as science communication material.

